# Using Deep Learning to Predict Human Decisions, and Cognitive Models to Explain Deep Learning Models

**DOI:** 10.1101/2021.01.13.426629

**Authors:** Matan Fintz, Margarita Osadchy, Uri Hertz

## Abstract

Deep neural networks (DNN) models have the potential to provide new insights in the study of human decision making, due to their high capacity and data-driven design. While these models may be able to go beyond theory-driven models in predicting human behaviour, their opaque nature limits their ability to explain how an operation is carried out. This explainability problem remains unresolved. Here we demonstrate the use of a DNN model as an exploratory tool to identify predictable and consistent human behaviour in value-based decision making beyond the scope of theory-driven models. We then propose using theory-driven models to characterise the operation of the DNN model. We trained a DNN model to predict human decisions in a four-armed bandit task. We found that this model was more accurate than a reinforcement-learning reward-oriented model geared towards choosing the most rewarding option. This disparity in accuracy was more pronounced during times when the expected reward from all options was similar, i.e., no unambiguous good option. To investigate this disparity, we introduced a reward-oblivious model, which was trained to predict human decisions without information about the rewards obtained from each option. This model captured decision-sequence patterns made by participants (e.g., a-b-c-d). In a series of experimental offline simulations of all models we found that the general model was in line with a reward-oriented model’s predictions when one option was clearly better than the others.

However, when options’ expected rewards were similar to each other, it was in-line with the reward-oblivious model’s pattern completion predictions. These results indicate the contribution of predictable but task-irrelevant decision patterns to human decisions, especially when task-relevant choices are not immediately apparent. Importantly, we demonstrate how theory-driven cognitive models can be used to characterise the operation of DNNs, making them a useful explanatory tool in scientific investigation.

**Author Summary:** Deep neural networks (DNN) models are an extremely useful tool across multiple domains, and specifically for performing tasks that mimic and predict human behaviour. However, due to their opaque nature and high level of complexity, their ability to explain human behaviour is limited. Here we used DNN models to uncover hitherto overlooked aspects of human decision making, i.e., their reliance on predictable patterns for exploration. For this purpose, we trained a DNN model to predict human choices in a decision-making task. We then characterised this data-driven model using explicit, theory-driven cognitive models, in a set of offline experimental simulations. This relationship between explicit and data-driven approaches, where high-capacity models are used to explore beyond the scope of established models and theory-driven models are used to explain and characterise these new grounds, make DNN models a powerful scientific tool.

## Introduction

The main objective of cognitive psychology is to understand and explain how people think, infer and process information, usually by observing their behaviour under experimental conditions. In the field of decision making, researchers try to evaluate how people make decisions, for example by describing the way information is processed [1–3], or by understanding how the value of options is learned and updated through experience [4,5]. Such approaches use computational models that explicitly implement theoretical assumptions about the processes at hand, evaluated in a carefully constructed experimental design. However, as these models make specific assumptions about human behaviour and motivations, they may fall short if people’s behaviour is carried out in a completely different manner. An alternative approach is to use high capacity data-driven models not constraint by prior assumptions, which has the potential of breaking new ground in cognitive psychology [6,7]. However, relying on data driven models entails a different problem – the explainability or interpretability problem [8,9]. The way data driven models such as deep neural networks (DNN) operate, i.e., how they draw predictions from data, is usually opaque, and they are sometimes referred to as Black-Box models; therefore, their ability to explain how an operation is carried out is limited [10]. The current work demonstrates the use of a data driven model the Long Short Term Memory (LSTM) network, which is a type of recurrent DNN, as an exploratory tool for identifying predictable and consistent human behaviour in decision making beyond the scope of theory driven models. After identification of such behavioural patterns, we demonstrate that theory driven models can be used to characterise and explain the performance of DNNs.

DNNs are becoming a popular tool in cognitive neuroscience [11]. DNNs have the capacity for implementing and representing complex computational steps, allowing them to reach human-like performance level in some tasks (and super human in others) without specifying an explicit description of the process [6,7,12]. This can be done by training a DNN model to perform a specific task, specifying the criterion for successful task performance, usually in the terms of accuracy or achievement of a goal, defined by a loss function. Importantly, the manner in which the task should be carried out is not specified. For example, a DNN model trained to perform a visual object identification task can shed light on the information processing carried out by different brain areas in the visual pathways [13]. However, a potential limitation of the above approach of matching a DNN to a task involves the criterion for successful performance of the task, as humans may perform a task with a very different goal from that assumed by the researcher and intended by experimenter. For example, when training a DNN to play Go, one may set the loss function as winning the game, making the model attain super human capabilities [14]. However, unlike human Go players, this model will not allow a child to win once in a while to encourage him or to build his confidence, failing to predict real life human behaviour [15]. To overcome this problem, we suggest training a model with the goal of predicting human behaviour in a task, rather than with the goal of performing well in the task.

We are especially interested in capturing predictable behavioural patterns that seem to have different objectives than that intended by the experimenter.

In value-based decision making problems such as the restless bandit [16], a player can acquire rewards by sampling actions on a trial by trial basis, while keeping track of the actions’ ever changing reward probabilities. In this task, it is natural to suggest training a model with the objective of acquiring as much rewards as possible and examining how it corresponds with participants’ behaviour. Indeed, many of the cognitive-process models used to capture behaviour in such a scenario, most notably reinforcement learning [17], assume that people’s behaviour in such tasks is driven by reward and by the maximisation of reward [16]. When people’s behaviour seems to digress from this reward-oriented prescription it is usually dismissed as noise, random explorations or failure to learn the task by a researcher [18].

However, there are some indications that reward maximisation is not the only parameter driving behaviour in such tasks [19]. People’s likelihood of choosing an action seems to increase simply because they chose it before, regardless of its outcome [19,20], and they follow fallacies such as the ‘hot-hand’ and ‘gambler’ fallacies, which assume that there is a pattern in what are actually random events (a signal in the noise) [21]. To expand our understanding of human decisions beyond reward-oriented behaviour, a high capacity DNN model can be a useful exploratory tool. To gain such an insight, it is important to train the model with the goal of predicting human decisions, rather than with the goal of gaining as much rewards as possible.

The strategy of training a DNN to perform a task or to predict human behaviour in that task with the goal of gaining insights on how the prediction is made, poses another problem – how to interpret the model’s operations. DNN models are opaque by nature, also known as ‘black box’ models [7,10,22]. As DNNs usually include many parameters and complicated architecture, and are trained without the control of an experimenter, they provide little information about the computations they carry out, limiting our ability to derive insights from DNNs [9]. This problem is not specific to the use of DNNs as a research tool, and is prevalent in the field of machine learning, known as the interpretability problem, and the effort to explain machine behaviour is known as the explainability problem [22]. This problem is especially restrictive when trying to use such models to gain insights about the operation of a system [7]. Several researchers used different approaches to overcome this problem. One approach was to train many different models with different goals, and examine how they perform in predicting human behaviour, thus controlling for the model’s goal [23], and another approach was to use adversarial examples that meant misleading a model and thus gaining insights on its operations [24]. We suggest another direction, which is to use tools from cognitive neuroscience, the same explicit cognitive models described above, to characterise and explain the operations of such a data-driven black box model.

Here we used high capacity DNN as an exploratory tool to uncover patterns of human decision making beyond those captured by reinforcement learning models, and explicit cognitive-process models to characterise these new behavioural patterns. To do so we trained a DNN model with the goal of predicting human decisions in a four-armed restless bandit task. We examined the difference in predictive abilities between the DNN and a reward-oriented reinforcement learning model. To characterise the added predictive power of the DNN model, which may capture behaviour that is not reward-oriented, we used another DNN model to predict human decisions but without providing it with any information on rewards. By using off-policy simulations of the general DNN, i.e., the reward-oriented model, and the reward-oblivious DNN, we characterised the different policies and behaviours captured by the general DNN model.

## Results

### Task

We examined a dataset of human decision making in a four-armed bandit task [25]. The experiment included 965 participants, who played 150 rounds of a four-armed bandit task online (Figure 1). The participants were instructed to choose between four doors by clicking the door’s number to maximise the overall reward. We refer to these decisions as actions *a*_*t*_ (or *action at time t)*. Behind each door was a reward that drifted through the rounds in values ranging from 1 to 98, denoted *r*_*t*_ (or *reward at time t)*. The rewards behind the four doors were predefined, and three such payoff structures were used, in line with previous work [16].

**Figure 1:**
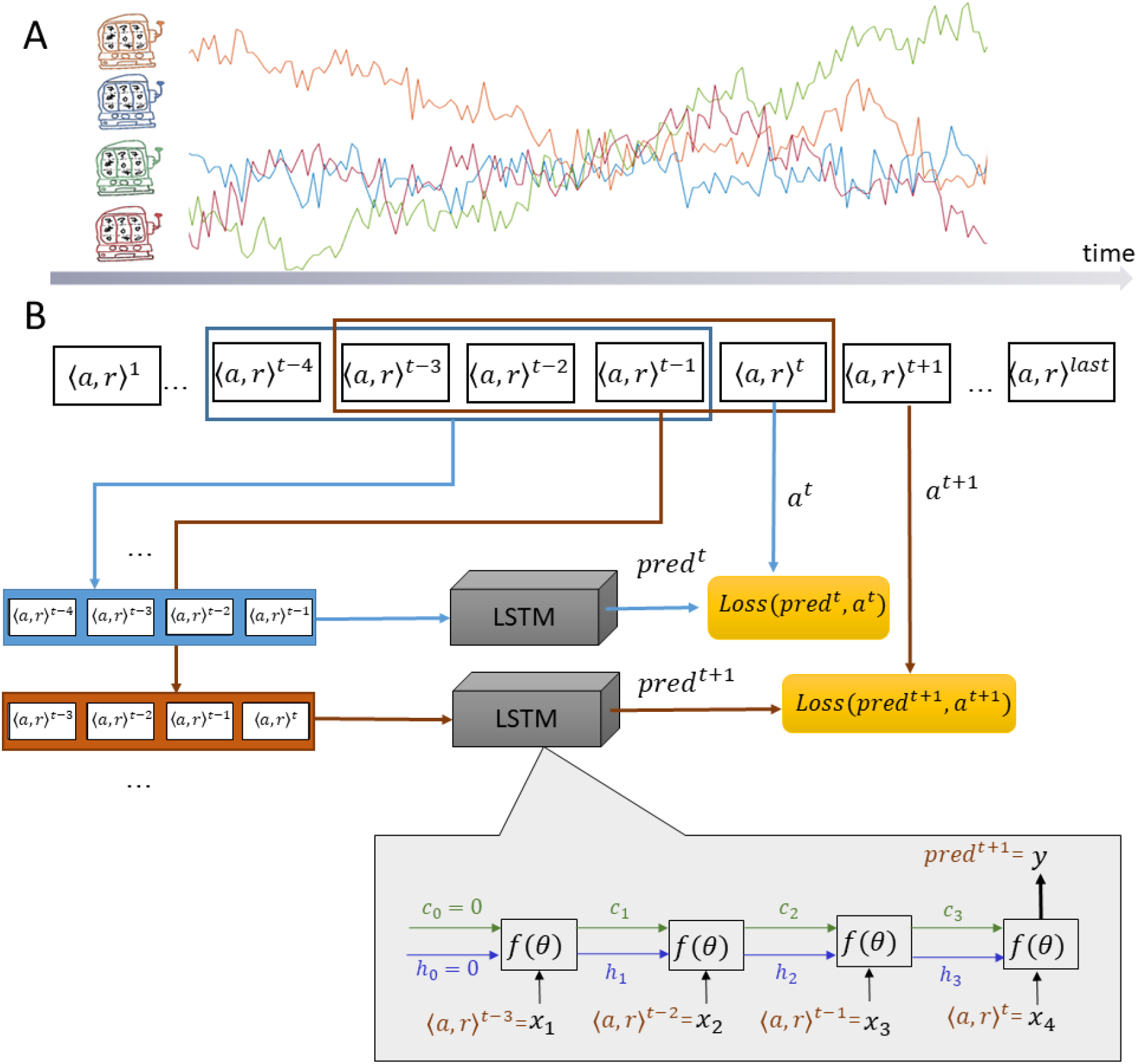
Experimental design and General LSTM model. (A) Experimental design – participants are asked to choose one of four options to gain rewards. The rewards associated with each option change slowly over time. (B) Sequences of four consecutive actions and rewards were used to train an LSTM model to predict the fifth action taken by the participant.

### General Model - LSTM

We used Long Short Term Memory (LSTM) to investigate human decision making in the above task. LSTM is a type of recurrent neural networks (RNN) that allows modelling temporal dynamic behaviour by incorporating feedback connections in their architecture (Figure 1). We trained the LSTM model to predict the participant’s action at time *t*, given his/her *K* previous actions and the corresponding rewards (in times *t* − *K*, . ., *t* − 1). We choose *K* = 4 as the number of action-reward steps to use in order to predict the next choice based on the experiments described in the supplementary materials. In sum, it was a trade-off between efficient use of the data and the memory needed for successful performance of the task, and using more than four action-reward pairs did not yield a significant improvement in accuracy. The LSTM model is trained on short sequences of action-reward pairs extracted from the participants’ data (as shown in Figure 1). The model can be viewed as a sequence of four units, corresponding to four consequent times. Each unit receives an input (denoted as *x*_*i, i*_ = 1, . . ,4 in Figure 1) comprising the participant’s action *a* and reward *r* from the previous step and the internal states (denoted *h*_*i*_,*c*_*i*_), which carry on the information from the previous 1, …, _*i*_ steps. Each unit is a complex function parameterised by a large set of parameters, but these are shared by all units. The parameters are learned from the training set by minimising the disparity between the model’s prediction at the fifth step and the corresponding action of the participant. The disparity is formulated as a cross-entropy loss over four outcomes.

Our goal was to capture different behavioural types of participants. Due to the large capacity of LSTM, we believe that it can learn different policies from a data set including many participants and generalise over participants that were not in the training set. To this end we performed a 5-fold cross validation in which the split was done over participants. Namely, each fold included 80% of the participants for training and 20% for testing. This way, instead of generalising the behaviour of the learned participant over time (as done in previous work [6]), we learn typical policies from one set of participants and generalise them over a new set of participants. The results of the cross-validation show that the trained LSTM model had an accuracy of 72.3%, indicating that it is capable of such generalisation.

### Reward-Oriented Model - Q-Learning

In order to examine how well the general model corresponded to a reward-oriented behaviour, i.e., actions that endeavour to maximise the acquired reward, we matched a reinforcement learning model to the data using a q-learning algorithm [17,26]. This model assumes that participants make decisions based on the learned value associated with each option. The value of each option is updated whenever the participant chooses this option and obtains a reward according to the prediction error, i.e., the difference between the obtained reward and the options’ expected reward (current value). A free parameter, learning rate, controls the amount of updating in each trial and another free parameter, inverse temperature, controls the stochastic nature of choices (how likely are participants to choose a low value option). These parameters were estimated for each participant individually, and were used to obtain model accuracy by comparing the model’s prediction of the participants’ choices.

This type of model is extensively used to model behaviour in such tasks [16,18,27,28]. Although many different extensions and elaborations of these simple mechanisms are used to capture different nuances in participants’ behaviour, they all share a common approach, which is that decisions are made to maximise reward, and that the history of obtained rewards drives the formation of reward expectations. These models are all reward-oriented in this sense.

### Comparison between the General Model and the Reward-Oriented Model

We found that the general model showed greater accuracy than the reward-oriented model in predicting participants’ actions in the task (General LSTM accuracy: 72.3%, Reward-oriented accuracy: 67.7%). Figure 2 compares the accuracy of the general model and of the reward-oriented model across participants. While the advantage of the general model was not very big for some participants, for others this disparity was substantial. The analysis revealed a subset of participants whose actions were not captured by the reward-oriented model, suggesting that their choices were not affected much by their actions’ outcome, but were nevertheless captured and accurately predicted by the high-capacity LSTM model.

**Figure 2:**
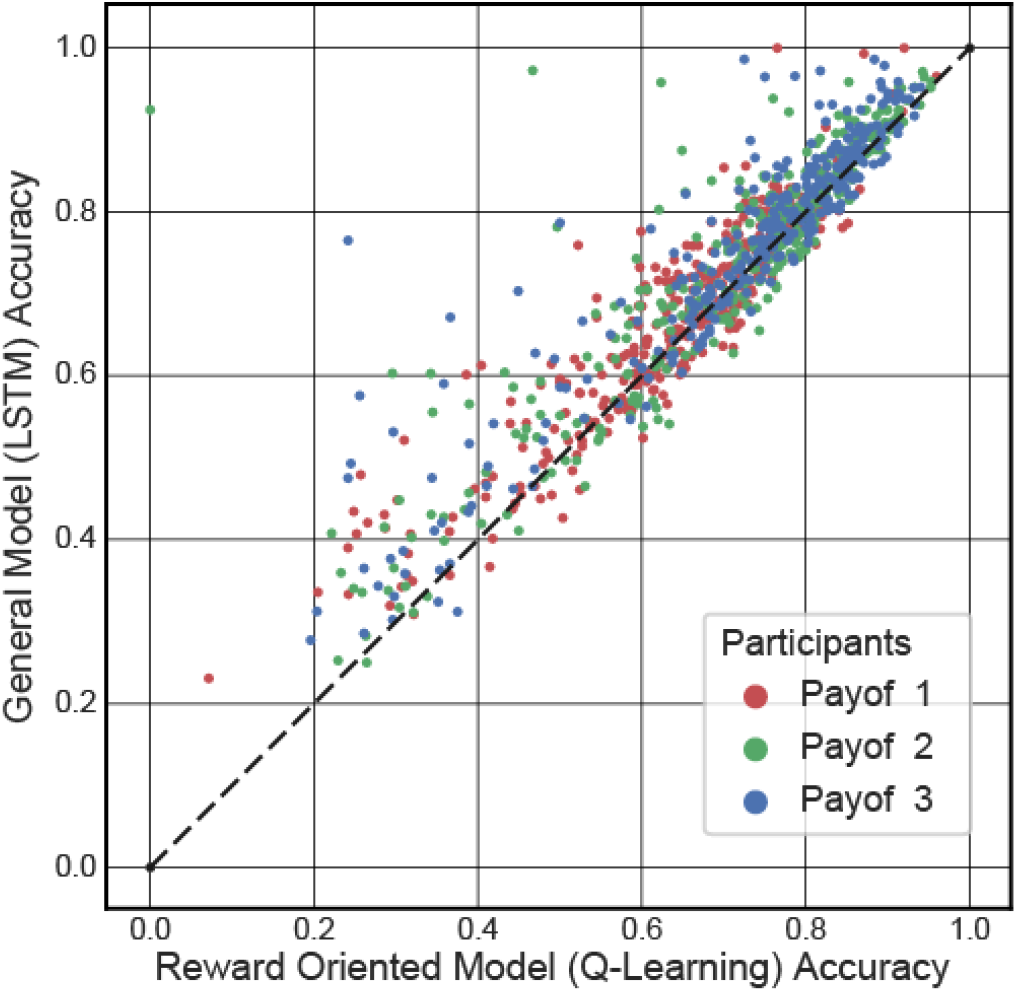
Comparison of the general and reward oriented models’ accuracy across participants. Different colours mark the different payoff structure the participants experienced. Dots in the top triangle represent participants whose actions were more accurately predicted by the general model than by the reward-oriented model.

We examined the models’ accuracy over time to identify periods during the task when the general model outperforms the reward-oriented model (Figure 3). In this analysis it was possible to observe that the disparity between the general model and the reward-oriented model varied over time and was most apparent during the period of uncertainty in the reward structure (see the accuracy of the models with respect to the reward structure in Figure 3 and in supplementary materials). Both models’ accuracy levels were high when one option was markedly better than others. However, the general model was more accurate when the options’ expected rewards were relatively close to each other. These observations were validated by applying the McNemar test of accuracies in each time point, showing a cluster of significant advantage for the general model in these periods. These times were associated with higher rates of exploratory choices, defined as times when participants choose options that are not associated with high reward [16].

**Figure 3:**
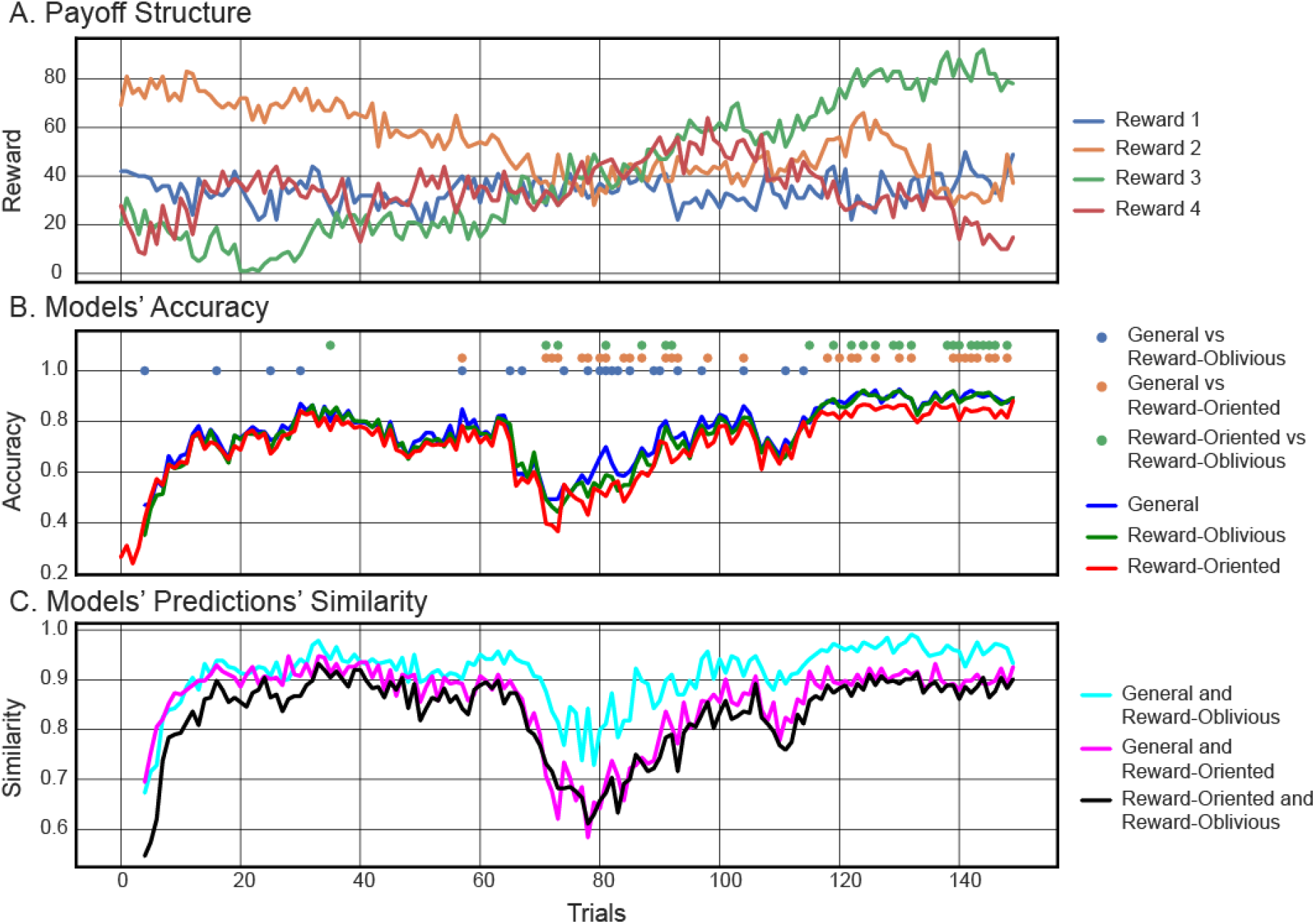
Analysis of models’ prediction over time. (A) Payoff structure indicates times when all options were similar and times when one option was distinctively better than others (payoff structure 1, for other payoff structure see supplementary materials). (B) Models’ prediction accuracy on trial-by-trial basis. Dots represent a p-value < 0.05 in the McNemar test between the two models’ accuracy rates. (C) Measure of similarity in predictions between the two models over time.

### Exploring the Disparity with the Reward-Oblivious Model

To better understand and characterise the added predictive capacity, i.e., what is it that the general model captures but the reward-oriented model fails to capture, we introduced a new, reward-oblivious model and compared the predictions of all models. We hypothesised that some participants may display behaviour that is not dependent on the action’s outcome, i.e., that is oblivious to reward [19,29,30]. One such pattern is the constant selection of the same action which is not very sensitive to the trial-by-trial fluctuations in outcome, for example when one option is much better than others [31]. During periods of uncertainty in outcome the actions are more diverse, which corresponds to exploration. However, participants may use specific patterns of exploration which are not reward dependent. For example, they may repeat sequences that were previously associated with outcomes [32], follow a pattern of exploring one option at a time or simply follow a motor pattern [33]. It is therefore possible that the gap between the general and reward-oriented models’ predictions could be captured by the reward-oblivious model.

To build the Reward-Oblivious model, we constructed a dataset from the original participants’ chains of actions, but without their rewards. We trained an LSTM model on this data by chopping it into 4-step action sequences and trained the model to predict the following action, similar to the general LSTM model. While actions, even stripped from rewards, may include information regarding the statistical regularities of rewards, this model is able to capture action patterns that are not reward oriented. Importantly, differences between the general LSTM and the reward-oblivious LSTM could be directly attributed to the explicit presentation (or lack) of actions’ outcome, as both models used the same architecture and training procedure. The results of 5-fold cross-validation showed that the reward-oblivious LSTM produces less accurate predictions of human behaviour than the general LSTM model (reward-oblivious accuracy: 69.9%). However, the reward-oblivious model is very good in pattern completion that is not reward-driven – it showed over 94% accuracy in predicting the next action for action sequences produced by off-policy simulations (detailed below).

We compare the average accuracy of all three models in predicting human behaviour over time in Figure 4. We observe that the reward-oblivious model is closer to the general model than the reward-oriented model in its accuracy. However, some discrepancy remains when there is no clear good option and the options’ outcomes are similar to each other, as validated by a McNemar test of accuracies. Predicting the next action for the constant pattern is easier than that of the exploration pattern (which is more diverse); this explains the better accuracy of the reward oblivious model when the best action is obvious (we believe that increasing the size of the training set could improve pattern prediction during uncertainty periods).

**Figure 4:**
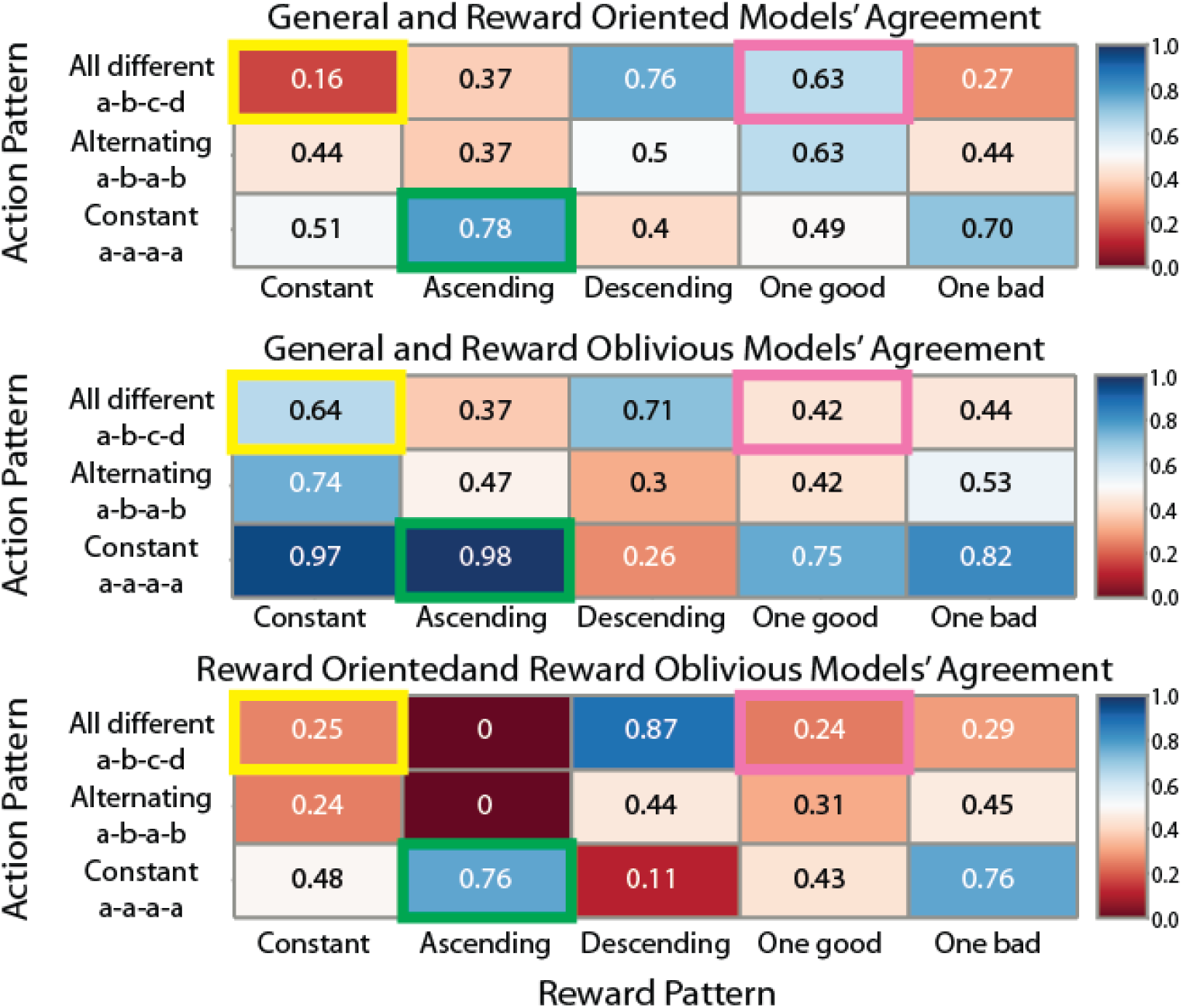
Comparison of models’ predictions in off-policy simulation. We simulated the models with different combinations of reward patterns and action patterns, and compared their predictions. We clustered together similar patterns in the different cells. Similarity values close to 1 indicate high similarity, and values close to 0 indicate low similarity.

To better understand the relationship between the reward-oblivious and the reward-oriented model predictions, we examined the overlap in their predictions over time (Figure 3-C). At each time point we compared their accuracy pattern - whether they were accurate or not in predicting each participant’s actions. If both models predicted correctly, we marked this mutual success, expressed as 1 in a similarity vector. If both failed, it was also marked 1 in the similarity vector. A mismatch in predictions was marked 0. Summing this similarity vector gave an indicator of similarity in predictions between the two models over time.

We observed that in many cases both reward-oriented and reward-oblivious models agreed, especially when one option was constantly better than others and therefore selected repeatedly (Figure 3). From a reward maximisation point of view, such a repeated choices pattern is the hallmark of exploitation - choosing the known best option [16]. From a reward-oblivious point of view, this was a pattern that regularly appeared in the choices sequences and therefore was relatively easy to capture and predict.

However, similarity in predictions decreased when there was no single obvious good option. As shown before, these were times where participants explored the different choices and the overall accuracy of all models in predicting participants’ choices decreased. The fact that models’ similarity decreased as well indicates that the models made different predictions in these times, suggesting that outcome information made a difference. Importantly, these were also times when the gap between the reward-oriented and the general model was the greatest, suggesting that maybe the advantage of the general model came from incorporating non-reward oriented choice patterns, which were captured by the reward-oblivious model, in order to form its predictions. Even though both the reward-oblivious and the reward-oriented model are less accurate than the general model in predicting human behaviour during periods of considerable uncertainty, they capture different aspects of human behaviour that jointly constitute the policy learned by the general model.

### Off-Policy Simulations

Examining the time course of models’ accuracy and similarity revealed that under some conditions the reward-oriented model and the reward-oblivious model gave very different predictions, and that both models may contribute to the performance of the general model. To better characterise these different predictions and their contribution we designed off-policy experimental simulations, where models’ predictions could be compared under specific action-reward sequences.

To start, we examined the predictions of the reward-oriented and reward-oblivious models for three action sequences coupled with two different reward sequences, resulting in six overall action-reward sequences (Table 1). We used a constant action pattern, i.e., choosing the same option over 4 consecutive trials, an all-different pattern where each option is selected once (1-2-3-4) and an alternating pattern where one alternates between two options (1-2-1-2). The rewards coupled with these action sequences either increased over time (10-30-50-70) or included one good outcome and three poor outcomes (40-80-40-40).

**Table 1:**
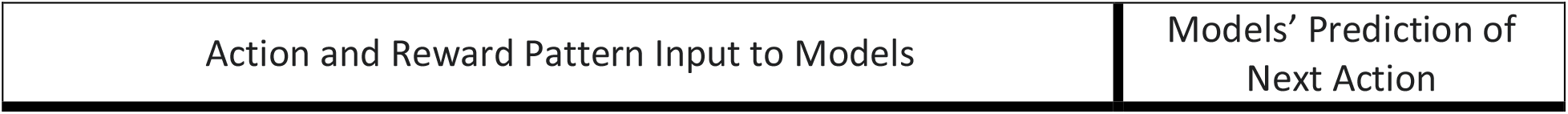
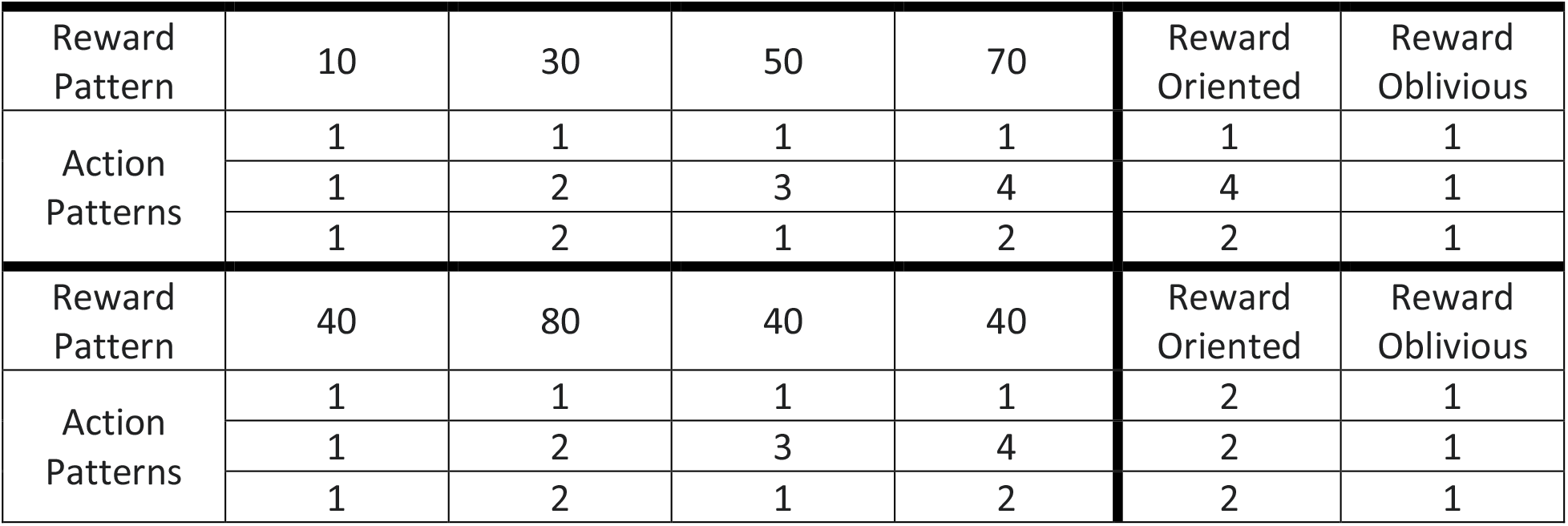
Comparison of the reward-oriented (q-learning) vs reward-oblivious models on action-reward examples. Q-learning predicts the action that maximises the expected reward, while the reward-oblivious model completes the pattern that exists in the sequence of previous actions.

| Action and Reward Pattern Input to Models | Models' Prediction of Next Action |
| --- | --- |

|  |  |  |  |  |  |  |
| --- | --- | --- | --- | --- | --- | --- |
| Reward Pattern | 10 | 30 | 50 | 70 | Reward Oriented | Reward Oblivious |
| Action Patterns | 1 | 1 | 1 | 1 | 1 | 1 |
|  | 1 | 2 | 3 | 4 | 4 | 1 |
|  | 1 | 2 | 1 | 2 | 2 | 1 |
| Reward Pattern | 40 | 80 | 40 | 40 | Reward Oriented | Reward Oblivious |
| Action Patterns | 1 | 1 | 1 | 1 | 2 | 1 |
|  | 1 | 2 | 3 | 4 | 2 | 1 |
|  | 1 | 2 | 1 | 2 | 2 | 1 |

The reward-oblivious model predicted completion or repetition of the pattern, regardless of rewards (Table 1). It predicted that the participant would keep on choosing the same option in the constant pattern. In the all-different and alternating patterns, it predicted that the pattern would repeat itself, i.e., the first choice in the 4-action sequence would be chosen again. For example, after observing a pattern of 1-2-3-4 it predicted that the participant would choose 1.

The reward-oriented model’s predictions were tightly linked to the rewards (Table 1). For example, when observing the action sequence 1-2-3-4 coupled with the reward sequence 40-80-40-40 it predicted choosing option 2, as this option was associated with the highest reward, but when the same action pattern was coupled with increased rewards, 10-30-50-70, a repetition of the last option, 4, was predicted, as it yielded the highest reward.

As these examples demonstrated, some action-reward sequences led to different predictions by the reward-oblivious and reward-oriented models. We expanded these sequences to cover a variety of action-reward sequences, falling into the same categories of action patterns (constant, alternating, all-different) and rewards (increasing, decreasing, constant, one-good, one-bad) (see methods and supplementary materials for full details of the sequences). In this wide array of action and reward sequences we observed that the reward-oblivious model always predicted pattern completion, similar to the results shown in Table 1, indicating that this model indeed captures motor patterns of actions (see supplementary materials for detailed predictions by all models).

In Figure 4 we present the pair-wise comparison of the models’ predictions for the different combinations of action and reward sequences. These were clustered together to provide an easy overall description of the similarity in predictions between the three models. Importantly, we compared the agreements between the general model, the reward-oriented model and the reward-oblivious models, to identify specific action-reward sequences where the general model follows the reward-oblivious pattern-completion predictions and when it converges with the reward-oriented reward maximisation predictions.

A number of patterns emerged from these comparisons. In some cases, all models give similar predictions, in others the general model’s predictions seem to converge with the predictions made by the reward-oriented model, while in yet others they converge with the reward-oblivious model’s predictions. Examining the combinations of reward and action sequences that underlie these cases can help characterise the operation of the general model in terms of sensitivity to action patterns and association with rewards.

First, all three models generated similar predictions when the constant action pattern (a-a-a-a) was accompanied by an increasing reward pattern (green cells in Figure 4). As demonstrated above (Table 1), in such cases the pattern completion prediction made by the reward-oblivious model that the same action would keep on being chosen, is the same as the reward maximisation prediction made by the reward-oriented model that the action with the highest experienced reward would be chosen. Note, that this agreement in prediction drops quickly when the action pattern is not constant – pattern completion and highest experienced reward are associated with different actions.

Next, we observed action-reward combinations when the general model generated predictions similar to the reward-oblivious model (pattern completion) and different from the reward-oriented model (highest reward) (yellow cells in Figure 4). This happened when the rewards were constant, and most notably when these constant rewards were accompanied by an all-different action pattern. The reward-oriented model predicted actions randomly when they had the same expected reward (q-value), and avoided options with constant low reward. The general model and the reward-oblivious model generated pattern completion predictions, predicting repetition of the first action in the sequence.

Finally, we identified combinations of reward and action sequences when the general model predictions were in line with the reward-oriented model, and less so with the reward-oblivious model (pink cells in Figure 4). When the rewards sequence included one very high reward and three lower rewards (e.g., 40-80-40-40), the general model usually converged with the reward-oriented model in predicting the choice of the highest expected reward action. This was in contrast to the reward-oblivious model that predicted choosing according to the actions pattern.

The observation of agreement between the general model and the reward-oriented and reward-oblivious models under our experimental simulated conditions demonstrated the contribution of reward and action pattern information to the general model’s predictions. Generally speaking, when the rewards were relatively high and stable, in the constant rewards and one-bad reward sequences, the general model tended to favour pattern completion and ignored fluctuations in rewards. When the rewards were less stable, especially when one reward was markedly better than others, the general model’s predictions were governed by the expected rewards, and action patterns had less bearing on its predictions. Importantly, these results suggest that both action-pattern repetition and reward maximisation strategies were encoded by the general model, and may contribute to its observed higher accuracy across time and participants.

## Discussion

In this paper we took advantage of DNN models’ great capacity and ability to capture regularities in data, and used them as exploratory tools for examining the scope of predictable human behaviour in a widely-used learning and decision making experimental paradigm. We observed that our general model gave more accurate predictions than a reward-oriented reinforcement learning model usually used to model behaviour in this task. This disparity was more pronounced when the reward uncertainty was high. After identifying this gap, we set out to characterise what made the general DNN model more accurate. Based on previous literature in human decision making, which suggested that humans follow some task-irrelevant regularities [29,30,32], we hypothesised that this model may capture behaviour that is not reward-oriented, and therefore trained another DNN model, the reward-oblivious model, that did not have direct access to the outcomes of actions in this task. The reward-oblivious model’s predictions were associated with action pattern completion – it was likely to predict actions that complete a sequence of actions. Using experimental simulations, we found that the general model converged with the reward-oriented model’s predictions under some experimental settings, and with the reward-oblivious model’s predictions in others. Specifically, the general model predicted pattern completion when all actions were expected to lead to similar rewards, i.e., there was no clear best option to choose. It demonstrated that during high uncertainty, when the best course of action is not clear, people explore options following a *predictable* pattern of action. This reliance on patterns of actions could not be identified using the standard reinforcement learning, reward-oriented approach, demonstrating the use of DNN models as exploratory tools in cognitive science.

DNNs are an extremely useful tool, performing a variety of tasks across multiple domains [10,34], and specifically performing tasks that mimic human behaviour [14,35,36]. It is therefore not surprising that these tools were adopted by cognitive neuroscientists, examining how such models represent information and the extent to which the operation of such models is similar to information representation and processing in the brain [11,13]. However, these models’ level of complexity, and their tendency to track regularities in the data in an unpredictable and opaque manner, means that while they may perform well, their explanatory power is limited [7,12,37]. Fortunately, DNNs can be a useful scientific tool in a number of ways, one of them being for exploratory investigations [38]. In a recent work, Dezfouli et al. (2019) showed that a DNN can capture meaningful differences in behaviour between groups of participants diagnosed with bipolar and unipolar disorders, and healthy controls. Moreover, the DNN model was able to capture irregularities in the clinical populations’ choice patterns, that could be used to inform future research. Here we demonstrated that a DNN model could uncover patterns of behaviour made by healthy populations that diverge from the reward-oriented prescription of the experimenters. DNNs can therefore expand researchers’ field of view, casting light on behavioural patterns so far treated as noise by theory-driven approaches.

However, after the first exploratory step is taken, a second explanatory step is needed. This problem is not unique to scientific exploration, but is encountered by many researchers and practitioners trying to provide explanations of how their black-box models operate, in order to gain better understanding of how they work, what they predict, and to promote trust in these algorithms [8,9]. This is referred to as the explainability problem. Different tools and approaches are being developed for this purpose, for example using visualisation to make linear regression models easy and quick to understand, and matching decision tree models to provide a systematic description of the model’s behaviour [39–42]. In cognitive neuroscience, another approach to this problem is to use behavioural experimental tools to explain the model’s behaviour [6,10]. One way to carry out this task is by examining the different experimental settings that make the model fail, known as adversarial examples, [24], which has a long tradition in cognitive psychology, from the use of visual illusions to study perception to the characterisation of biases in decision making [43]. Another method is to train many models with different goals, and to examine which models best describe human behaviour [23]. Here we used cognitive models that provide explicit predictions, reward-oriented and reward-oblivious models, to characterise the performance of our general DNN. Using off-policy experiments designed to differentiate between these two models, we were able to chart the gap between the general DNN model and the traditional reward-oriented approach. Explicit cognitive models can therefore be useful for explaining the operation of DNNs, and for guiding future investigations following the exploratory use of DNN models.

To conclude, our work demonstrated how DNN models can be used to uncover hitherto ignored human behaviours. DNNs are suggested to be a useful exploratory tool in cognitive neuroscience, along with explicit, theory-driven models. The explicit models provide explanations of cognitive processes, and can be used to characterise data-driven DNNs. DNNs can expand the scope of operation for explicit models, leading investigators to examine behaviour previously considered noisy, by showing that in fact it predictably emerges from the data. This relationship between explicit and data-driven approaches can be useful for explaining the operations of black-box models outside the field of cognitive neuroscience, as the interaction between humans and such artificial models becomes more and more prevalent.

## Methods

### Task

We examined a dataset of human decision making in a four-armed bandit task [25]. The experiment was carried out online, and included 965 participants playing 150 rounds of a four-armed bandit task. In this task participants had to choose one of four options in each trial, in order to obtain rewards (points in the game, no performance based monetary reward was given in this experiment) (Figure 1). We refer to these choices as *a*_*t*_ (or *choice at time t*). The amount of rewards associated with each option was initially set to a value between 0 and 98 points, and drifted over time (standard deviation σ = 2.8), so participants had to keep on tracking the outcomes to pick the highest paying option. Rewards are denoted by *r*_*t*_ (*or reward at time t*). Participants faced one of three payoff structures generated in this manner (see Figure 1 and supplementary materials). Participants had to reach a decision within four seconds, and failing to do so moved them to the next trial with no reward. Of 965 participants in the dataset, only 127 players completed all 150 rounds, while others missed at least one trial. The average number of rounds per participant was ∼145. The average reward in each round across all participants was 65.8.

### Data preparation for models

For the general model, we created a data set by splitting the original data of each participant into 4-step action-reward sequences < (*a*_*t*−4_, *r*_*t*−1_), (*a*_*t*−3_, *r*_*t*−3_), (*a*_*t*−2_, *r*_*t*−2_), (*a*_*t*−1_, *r*_*t*−1_) > using a sliding window (Figure 1). After evaluating the gain in accuracy resulting from the addition of each step (see Figure SF1 in the supplementary materials), we chose 4-step sequences as a tradeoff between efficient use of the data and the memory needed for successful performance in the task. The model makes the prediction at time *t* given the previous 4 steps, thus the first prediction is for the 5^th^ round.

We excluded missed trials from the sequences, causing 4% of the sequences to include a gap (i.e., missed response), for instance a sequence where action 6 was missing can be < (*a*_4_, *r*_4_), (*a*_5_, *r*_5_), (*a*_7_, *r*_7_), (*a*_8_, *r*_8_) >, with the model predicting action 9. Almost all of the sequences with a gap missed only a single round. Since the drift in rewards associated with the options was slow, we included sequences with gaps in the dataset. The order of the resulting sequences across all participants was shuffled (as we assume no continuity beyond the 4-step sequence).

The reward-oblivious model inputs a sequence of actions without the corresponding rewards < (*a*_*t*−4_), (*a*_*t*−3_), (*a*_*t*−2_), (*a*_*t*−1_) >. The sequences were produced using the same procedure as for the general model.

The reward-oriented model was matched to the total action sequences of the participants (i.e., 150 actions and rewards), and was matched for each participant independently. To preserve the trial number of each action we marked missed actions with a value of ‘-1’, which indicated to the model to skip these actions.

### LSTM Models

Long Short Term Memory (LSTM) is a type of recurrent neural networks (RNN), which allows modelling temporal dynamic behaviour by incorporating feedback connections in their architecture [44]. LSTM networks have an internal mechanism called gates that can regulate the flow of information and yield improved performance compared to vanilla RNN. Each LSTM unit includes a **memory cell *c*** (Eq. 5) and three gates: an **input gate *i*** (Eq. 1), an **output gate *o*** (Eq. 2) and a **forget gate *f*** (Eq. 3), that operate on the flow of information arriving in the cell via input variable *x*^<*t*>^and feedback variable *h*^<*t*−1>^. The architecture of the LSTM unit is shown in Figure 6 and the equations for the forward pass are summarised in Eq. 4–6.

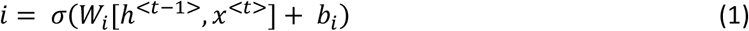

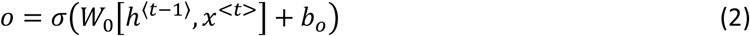

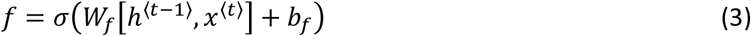

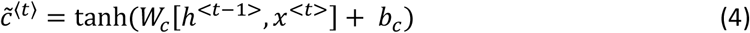

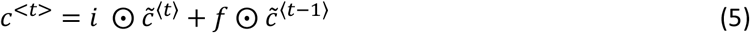

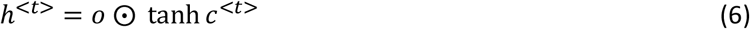

**Figure 6:**
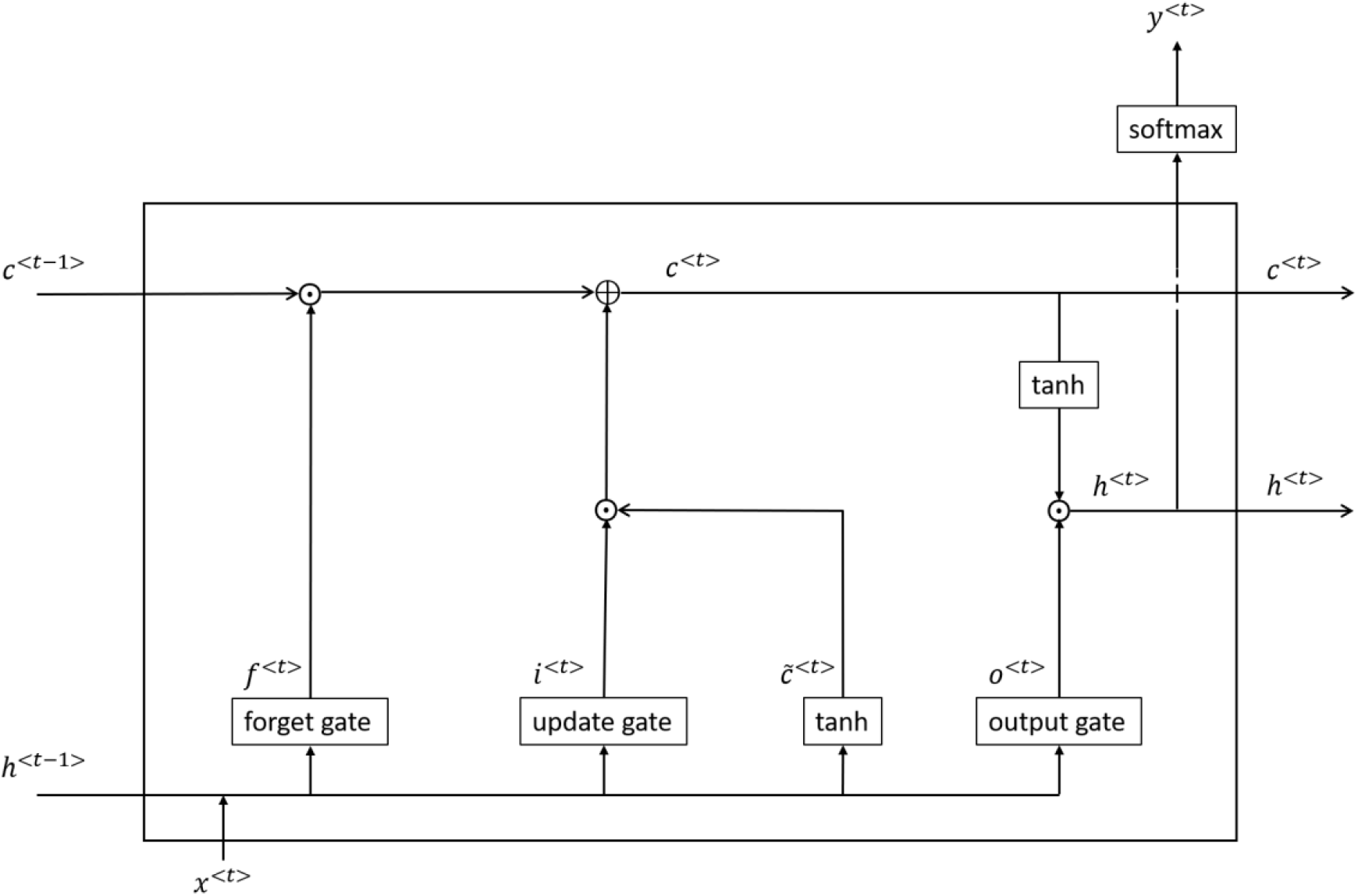
A LSTM cell. 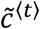 is a candidate for replacing the current cell state using previous feedback and current input [Eq. 4]. The new cell state *c*^⟨*t*⟩^ uses the forget and input gate (based on the previous step) along with the current candidate cell state to determine whether the cell state is to be updated or not. The next cell will get the final feedback *h*^<*t*>^and if it is the last cell in the sequence, a 4-way softmax receives the feedback and makes a prediction for the current time t.

Our general and reward-oblivious models are both LSTM models with four units. The last unit includes a 4-way softmax layer for outputting the probability of choosing each of the 4 doors. For the general model, the input *x*^<*t*>^ to the unit *t* comprises the previous action and previous reward (*a*_*t*−1_, *r*_*t*−1_), while the reward-oblivious model inputs only the previous action (*a*_*t*−1_).

The general and reward-oblivious models were trained on the general and reward-oblivious data, respectively using the Categorical Cross-Entropy Loss:

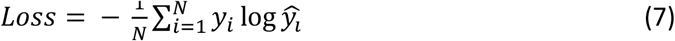

We performed a 5-fold cross validation. The evaluation phase was performed on the hold-out set in each fold without sliding window.

We used a single layer LSTM, as the task itself is simple and there is no need for a deeper network. Each LSTM cell has 64 hidden units, trained for 300 epochs with a batch size of 2048.

Both the hidden layer size and the number of epochs were determined after running a grid search for hyperparameter tuning.

The models were trained using Adam optimiser with a learning rate of 0.001, beta1=0.9 and beta2=0.99. We implemented the LSTM models in TensorFlow, under the Windows operating system, with the GTX 970 graphic card.

### Q-Learning Model

For the reward-oriented model we used a reinforcement learning model, q-learning [17,45], which is commonly used to model the behaviour of human participants in similar tasks to the one used here [16]. The model assumes that decisions are driven by the expected reward of each option, and that these expected rewards are learned on a trial-by-trial basis by updating the learner’s expectations (known as q values) based on prediction errors, the difference between the obtained reward and the expected reward. In our case, in each trial the participant makes a decision which of the four doors to open, noted as action a_t_, and receives a reward r_t_. In each round the expected value of the chosen action, noted as *Q*_*t*_(*a*_*t*_) is updated according to Eq. 8:

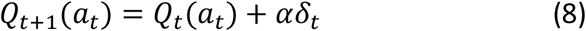

where 0 ≤ α ≤ 1 is a free learning rate parameter and *δ*_*t*_ is the prediction error *δ*_*t*_ = *r*_*t*_ − *Q*_*t*_(*a*_*t*_).

In each trial t, the model assumes that the participants make their choices according to a softmax distribution based on the q-values they learned so far:

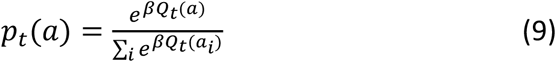

where 0 < *β* is a free parameter representing inverse temperature, i.e., the level of noise in decisions. This decision rule attributes higher probabilities to actions that are expected to yield high rewards.

Unlike our LSTM models that operate on sequences of four actions-rewards, the q-learning model is affected by the aggregation of the entire history of actions and rewards until the time point. This aggregation is shaped by the learning rate, such that the effect of previous actions and outcomes exponentially decays. However, unlike the LSTM model that keeps track of the ordering, in q-learning the specific order of actions and rewards is lost in the aggregation process.

The model was optimised for each participant’s entire sequence of actions and rewards, by maximising the log likelihood of actions (aggregated log probabilities of observed actions) using Eq. 8 and Eq. 9, with SciPy’s optimisation package (using L-BFGS-B). The optimisation process yielded an estimate of parameters *α* and *β* for each participant. This process also allowed recovery of trial-by trial Q and p(action) values for each of the four actions.

To calculate the accuracy of the reward-oriented model we chose the highest q-value in each time point as the model’s prediction, and compared it to the participant’s actual choice. The model’s accuracy was not calculated for trials where the participant did not make a choice.

### Model Comparison

To statistically evaluate the differences between the models’ accuracies in each trial, we used the McNemar test [45,46]. We calculated a contingency table for two models in each trial, summarising the number of correct predictions and incorrect predictions across participants. This process allowed us to identify quantitatively trials and periods within each payoff structure where models’ accuracies differ from each other. Comparisons were made in a pairwise manner and were not corrected for multiple comparisons as they were used to provide an estimation of the difference between the models and not for hypothesis testing.

To measure the similarity of models’ performance over time (Figure 3C), we compared the pattern of accuracy across participants for each trial, i.e., whether the models were accurate or inaccurate in predicting the same participants’ actions:

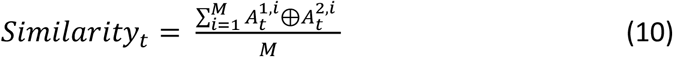

where 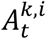 is the accuracy of the model’s prediction at time *t* for participant *i* = 1, . ., *M* for model *k* = 1,2 and ⊕ is XOR operation.

### Off-Policy simulation

To identify how reward-oriented and reward-oblivious predictions differ, and how they can be used to characterise the operation of the general model, we used experimental off-policy simulation. We used the LSTM models trained on the participants’ data as described above to generate action predictions for the general and reward-oblivious models. To simulate the reward-oriented model we used a q-learning algorithm with parameters set to the average of the individual estimated parameters.

The simulations included 3 types of action patterns: Constant (a-a-a-a), Alternating (a-b-a-b) and All-different (a-b-c-d). Five types of reward patterns were used: Constant (r1=r2=r3=r4), Ascending reward (r1<r2<r3<r4), Descending reward (r1>r2>r3>r4), One-good reward (e.g., r1>r2=r3=r4), and One-bad reward (e.g., r1>r2=r3=r4). We varied the exact rewards in each reward type and their timing in the one-different patterns. We also varied the specific actions used in the action patterns. An example of such action-reward combination is in Table 1. This resulted in 3400 combinations (136 reward sequences and 25 action sequences) of action and reward patterns overall, which we clustered into 15 actions and reward type combinations.

Models’ predictions were calculated for each of the action-reward combinations (see Table 1 for example), and were scored for agreement (1 = made the same action prediction, 0 = made a different prediction). Pairwise comparison of the model under the different action-reward types is presented in Figure 4. A link to all the model’s simulations is available in the supplementary material.

## Author contributions

M.F., M.O. and U.H. were involved in conceptualisation of the study and in the design of the analyses. M.F. carried out data analysis. M.F., M.O. and U.H. wrote the paper.

## Acknowledgments

U.H. and M.O. were supported by a University of Haifa Data Science Research Center (DSRC) seed grant. U.H. was supported by the Israel Science Foundation (1532/20).

