## Supplementary Figures for "Using Deep Learning to Predict Human Decisions, and Cognitive Models to Explain Deep Learning Models" (from another version of this preprint: version 1, 2021-01-15)

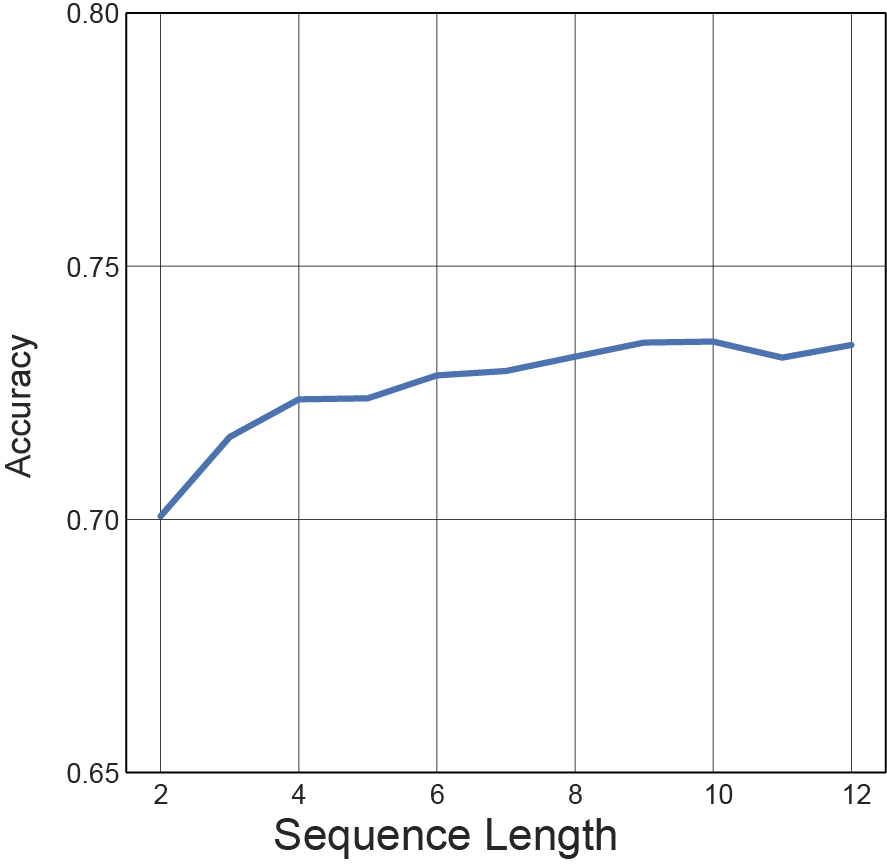

**SF1:** Accuracy of LSTM models with different sequence lengths. We trained LSTM models using different action-outcome sequence lengths, ranging from 2-12. The graph depicts the accuracy of each model in predicting participants’ actions. After evaluating the gain in accuracy made by the addition of each step, we chose 4-step sequences as a tradeoff between efficient use of the data and the memory needed for successful performance of the task.

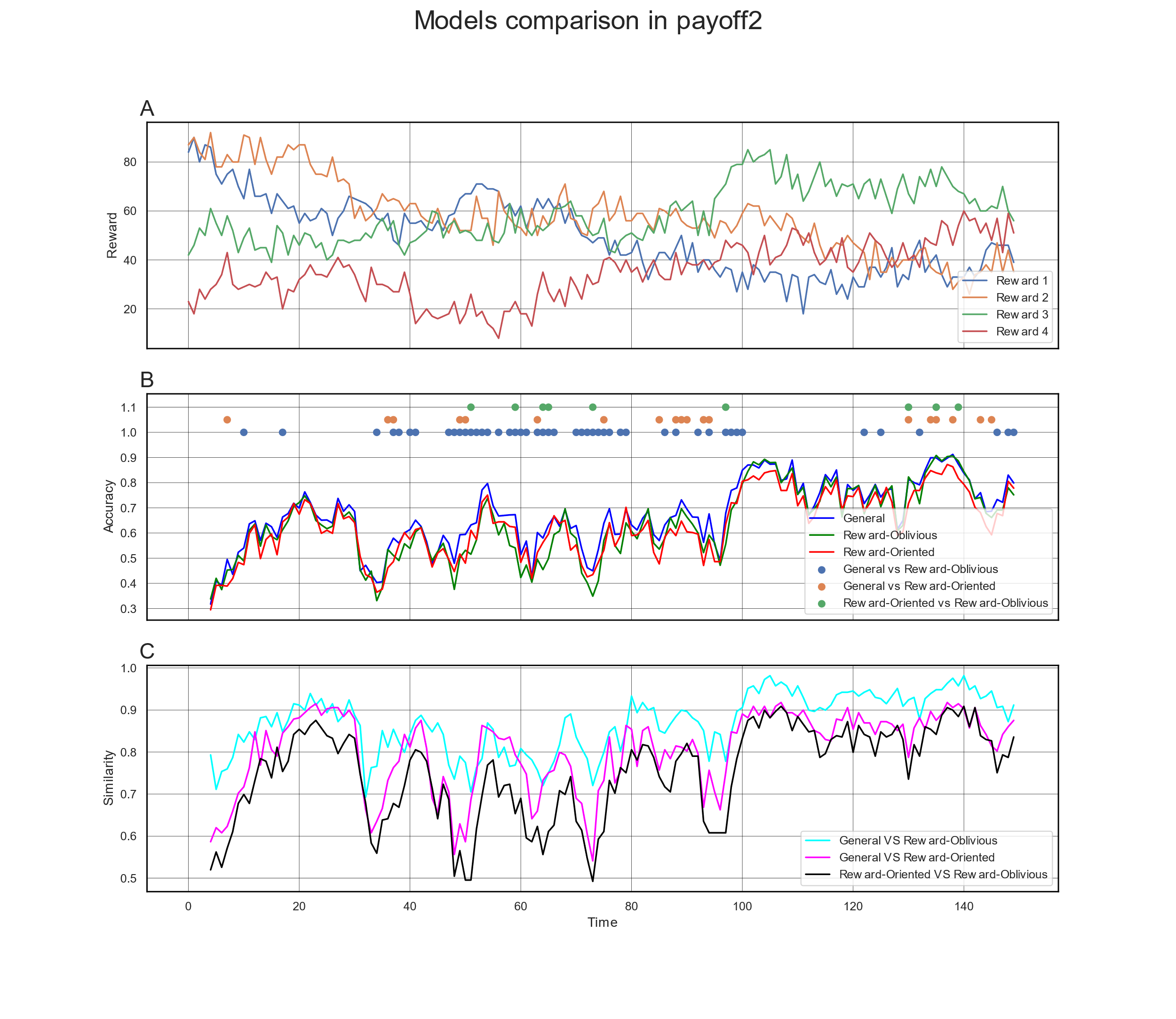

**SF2:** Analysis of models’ prediction over time. (A) Payoff structure indicates times when all options were similar and times when one option was distinctively better than in payoff #2. (B) Models’ prediction accuracy on trial-by-trial basis. Dots represent a p-value<0.05 in the McNemar test between the two models’ accuracy rates. (C) Measure of similarity in predictions between the two models over time.

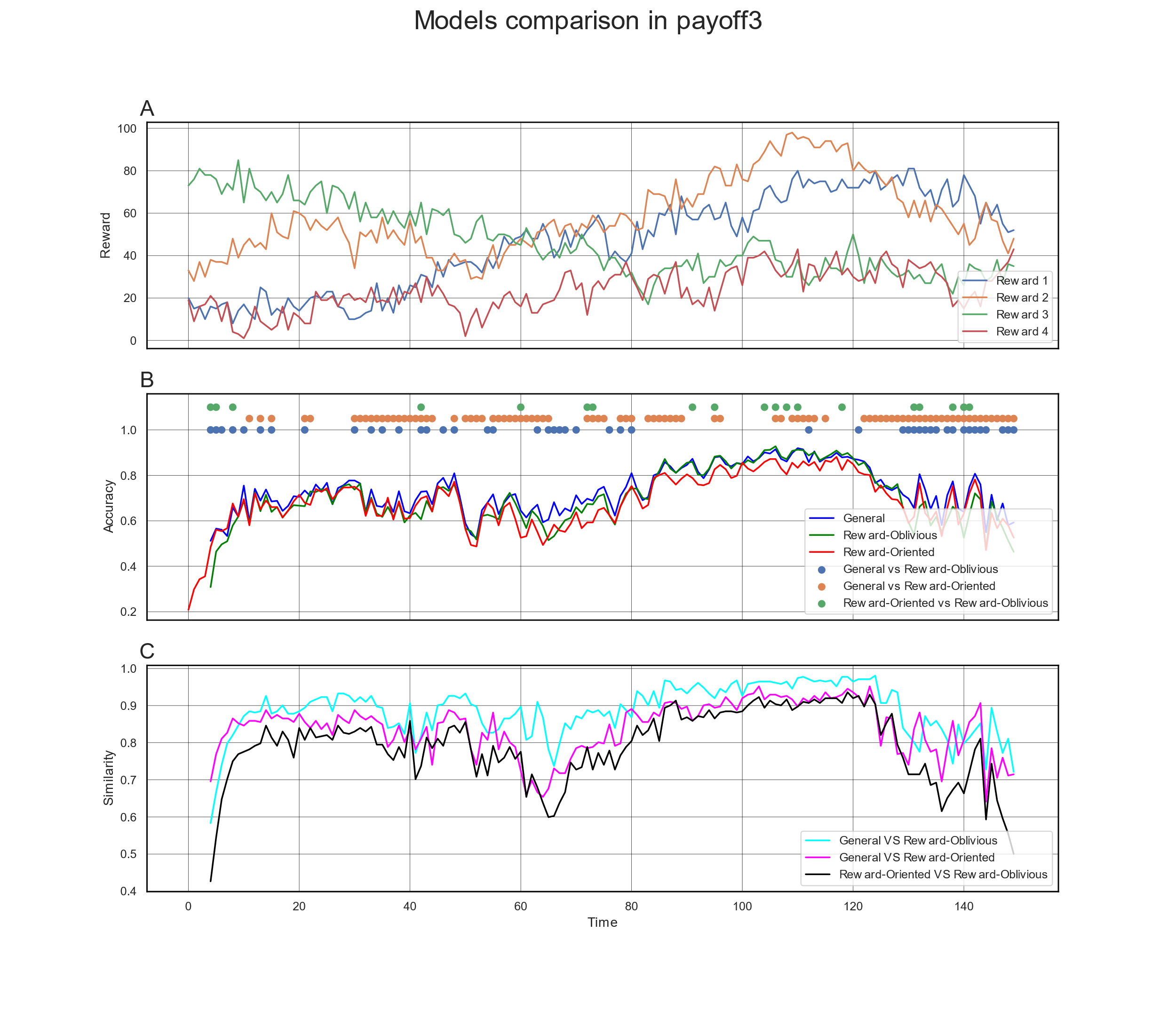

**SF3:** Analysis of models’ prediction over time. (A) Payoff structure indicates times when all options were similar and times when one option was distinctively better than in payoff #3. (B) Models’ prediction accuracy on trial-by-trial basis. Dots represent a p-value<0.05 in the McNemar test between the two models’ accuracy rates. (C) Measure of similarity in predictions between the two models over time.

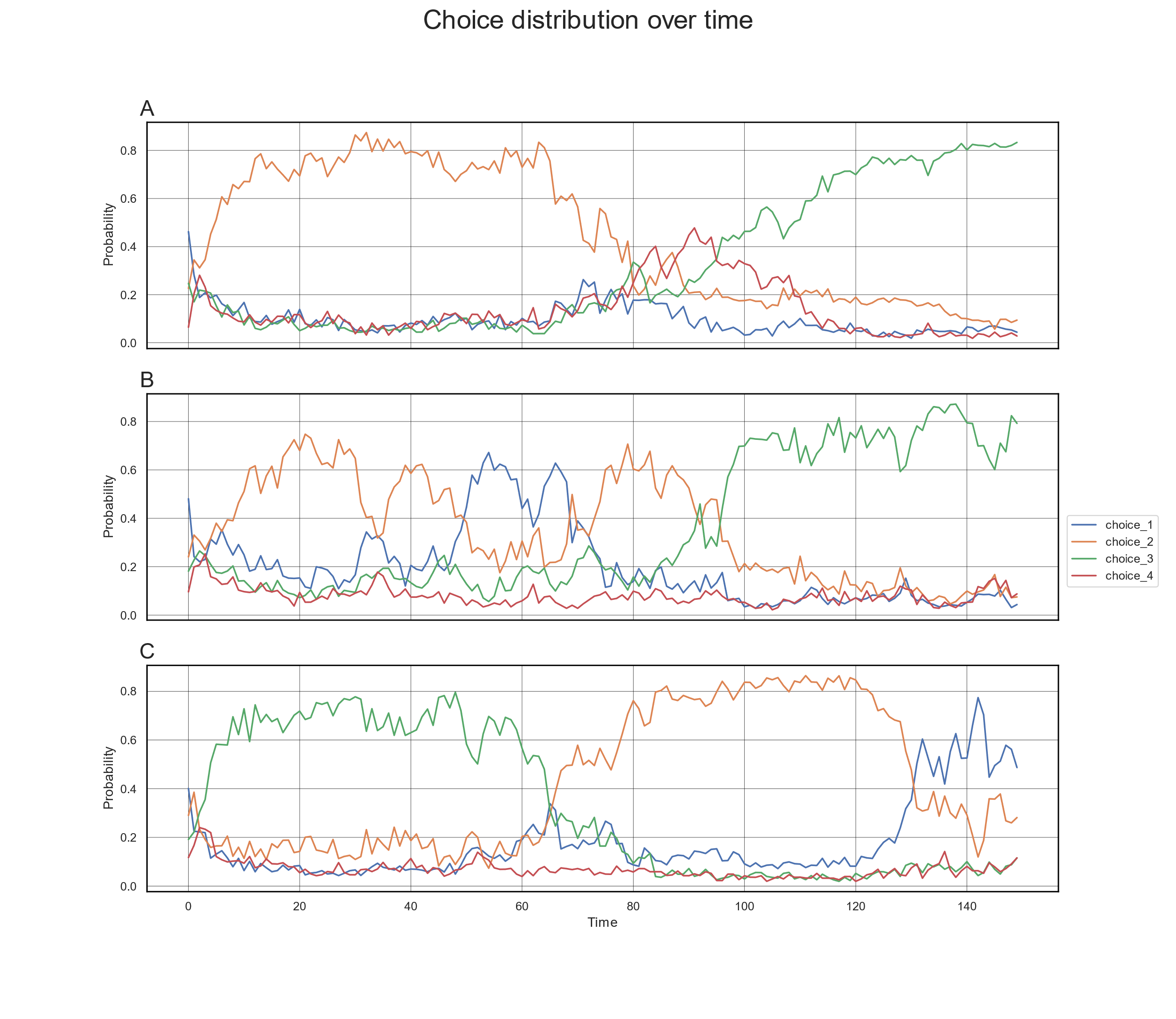

**SF4:** Distribution of actual choices by participants over time. (A) Payoff structure #1	(B) Payoff structure #2 (C) Payoff structure #3.

**SF4**

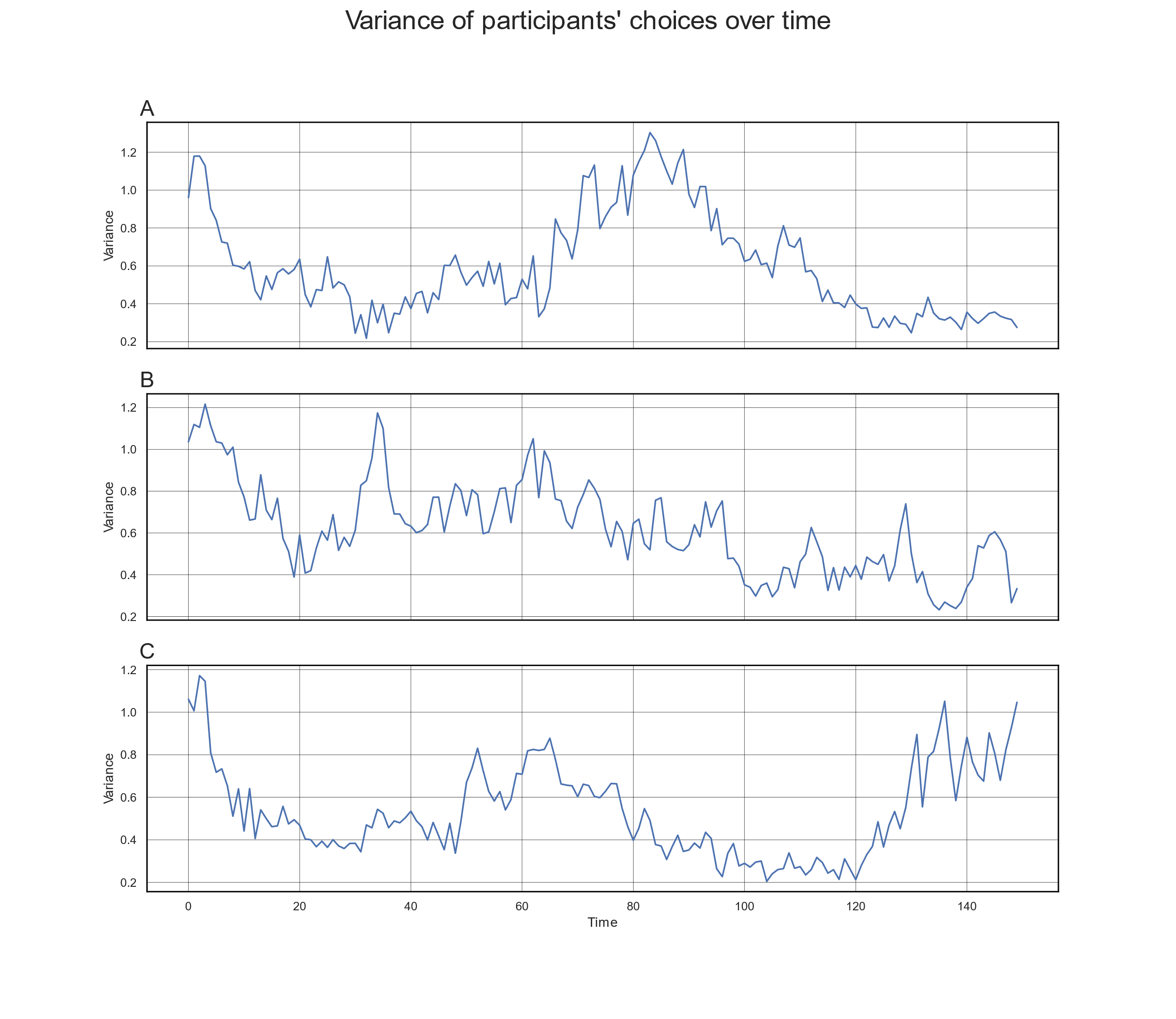

**SF5:** Variance of participants’ choices over time. (A) Payoff structure #1 (B) Payoff structure #2 (C) Payoff structure #3.

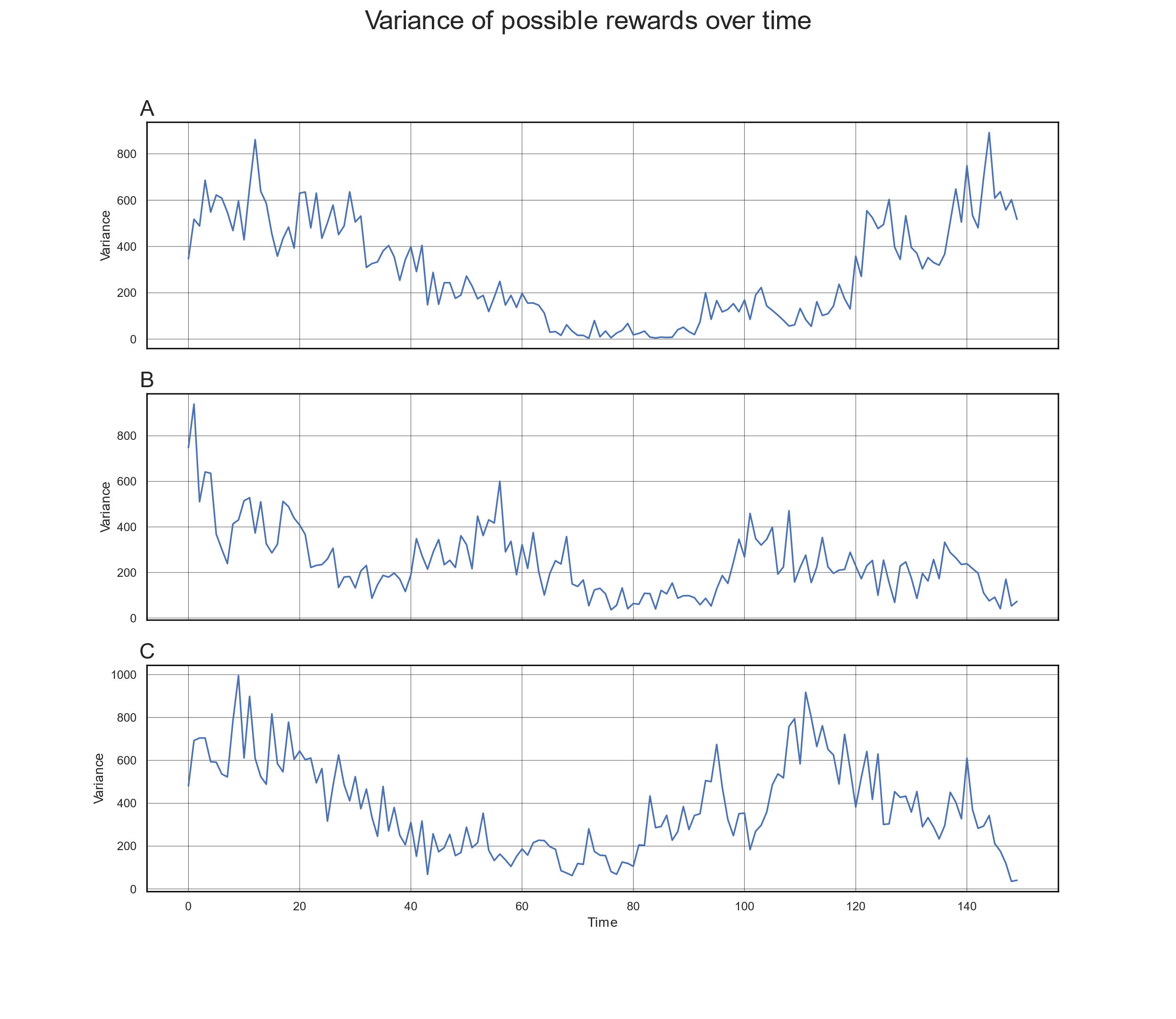

**SF6:** Variance of possible rewards over time. (A) Payoff structure #1 (B) Payoff structure #2 (C) Payoff structure #3.

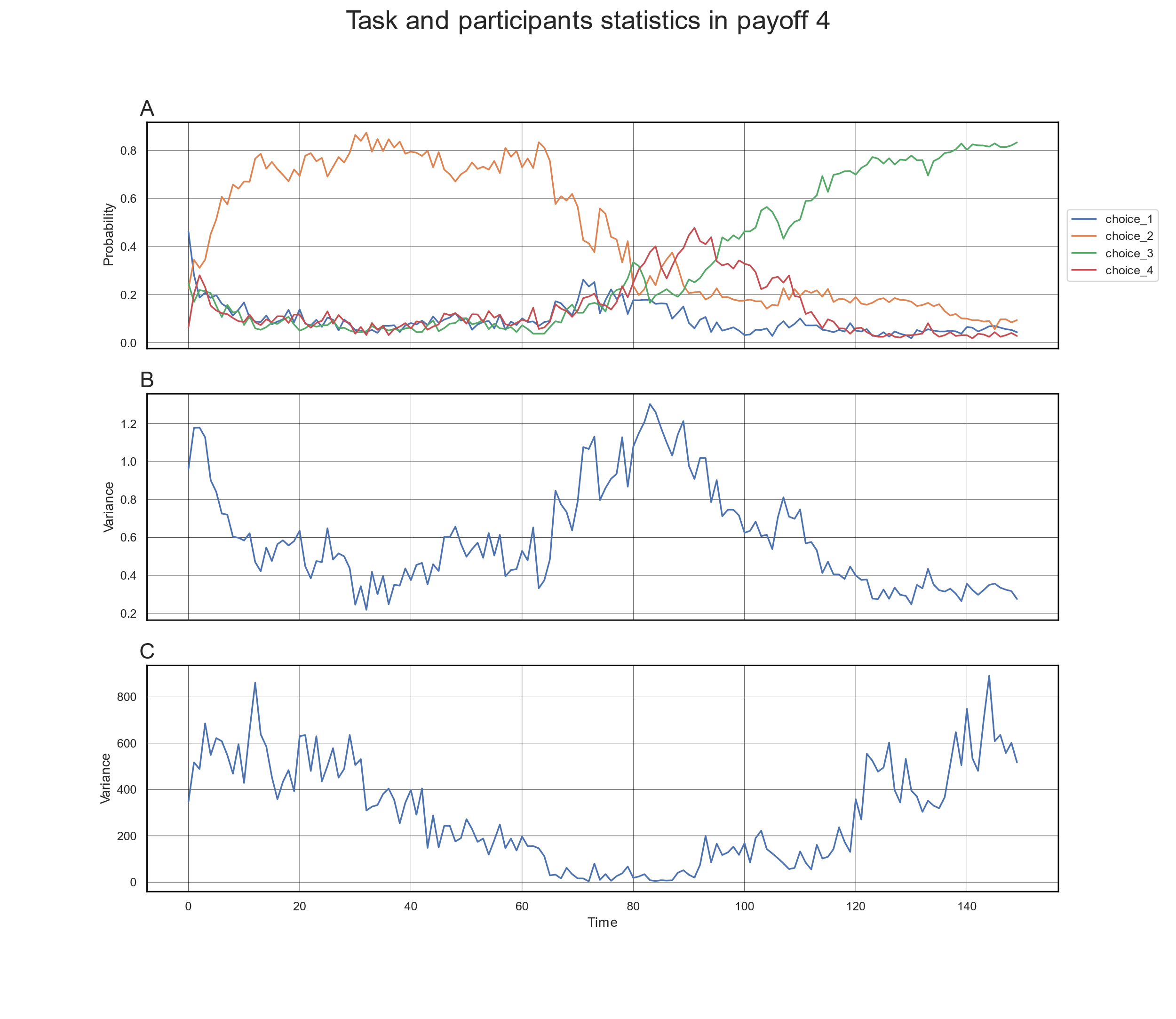
 **SF7:** Participants’ performance in the original task with payoff structure #1. (A) Distribution of actual choices by participants over time (B) Variance of participants’ choices over time (C) Variance of possible rewards over time.

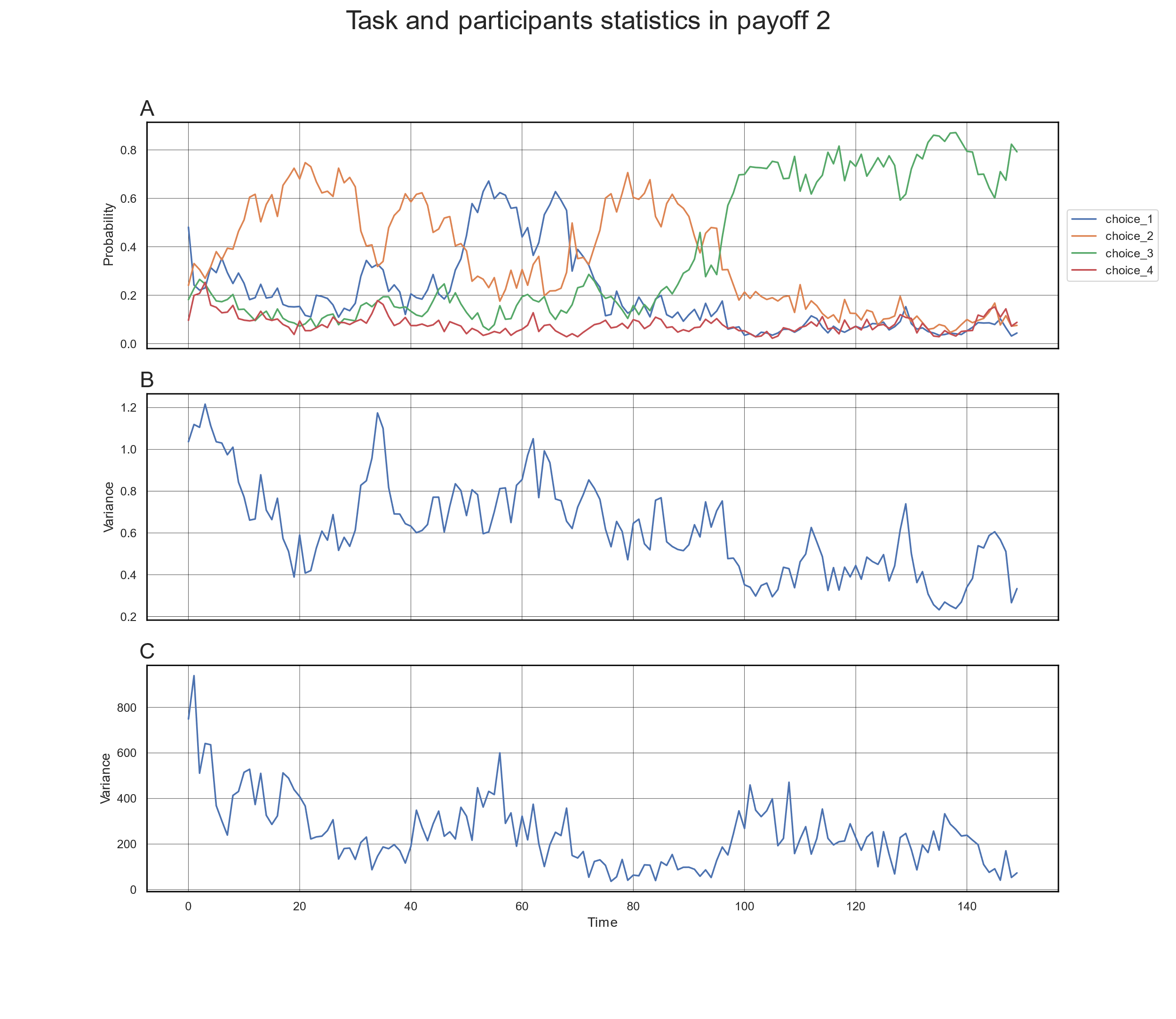

**SF8:** Participants’ performance in the original task with payoff structure #2. (A) Distribution of actual choices by participants over time (B) Variance of participants’ choices over time (C) Variance of possible rewards over time.

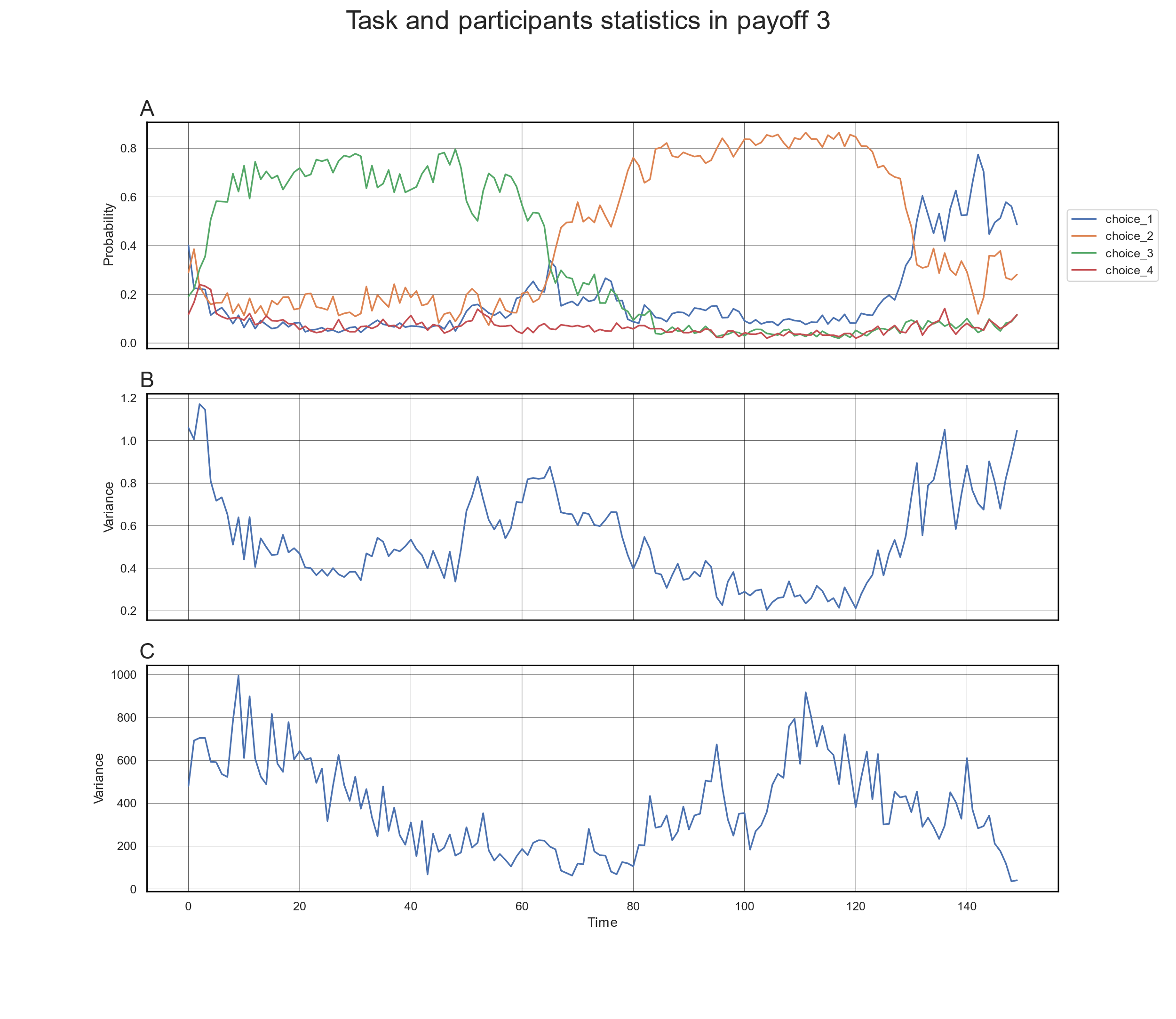

**SF9:** Participants’ performance in the original task with payoff structure #3. (A) Distribution of actual choices by participants over time (B) Variance of participants’ choices over time (C) Variance of possible rewards over time.
